# Integrating phylogenetic and network approaches to study gene family evolution: the case of the AGAMOUS family of floral genes

**DOI:** 10.1101/195669

**Authors:** Daniel S. Carvalho, James C. Schnable, Ana Maria R. Almeida

## Abstract

The study of gene family evolution has benefited from the use of phylogenetic tools, which can greatly inform studies of both relationships within gene families and functional divergence. Here, we propose the use of a network-based approach that in combination with phylogenetic methods can provide additional support for models of gene family evolution. We dissect the contributions of each method to the improved understanding of relationships and functions within the well-characterized family of AGAMOUS floral development genes. The results obtained with the two methods largely agreed with one another. In particular, we show how network approaches can provide improved interpretations of branches with low support in a conventional gene tree. The network approach used here may also better reflect known and suspected patterns of functional divergence relative to phylogenetic methods. Overall, we believe that the combined use of phylogenetic and network tools provide more robust assessments of gene family evolution.

## 1 Introduction

Advances in sequencing technology have lead to dramatic expansions in the number of sequenced genes within most gene families, both through the use of whole genome or whole transcriptome sequencing, or through broader taxon sampling. Gene families are generally studied through the use of phylogenetic approaches in order to identify closely and distantly related sequences, as well as to classify divergence between gene copies into those resulting from speciation (orthology) or gene duplication (paralogy)[1, 2]. Thus, phylogenetic approaches are widely employed to study how sequence divergence can lead to divergence of structure and/or function [3, 4]. When coupled with genome-context information, this approach can provide insightful understanding of gene regulation and function.

For instance, it is well-known that orthologous genes conserved at syntenic locations in the genome are more likely to exhibit conserved regulation [5] and function [6] than genes at nonsyntenic locations. On the other hand, the prevalence of whole genome duplications in plants poses challenges to the study of gene family evolution using exclusively phylogeny-based methods [3] due to the diverse outcomes of duplicated genes. Whole genome duplications produce syntenic paralogs that can be reciprocally lost [7, 8], sub or neofunctionalized [9], or even retained in the same functional roles as a result of relative or absolute dosage constraints [10].

A fundamental assumption of any phylogenetic reconstruction is that the observed changes occur exclusively through a hierarchical bifurcated branching process. This model is certainly a good representation of a major evolutionary force (i.e., descent with modification) [11] however many will argue that it fail to capture the diversity of evolutionary processes which shape the gene content of extant species [12, 13].

One way to address the complexity of evolutionary processes is to apply network approaches to address questions related to cell organization and functioning [14], human diseases relationships [15] and plant gene function prediction [16]. Network approaches have also been successfully applied to study fungi evolution based on enzymes related to the chitin synthase pathway [17]. Recently, Carvalho et al [18] have used a network-based approach to address the origin of the mitochondria, providing a new perspective on the study mitochondrial evolution.

Network-based approaches can overcome some of the limitations of phylogenetic methods. For instance, these approaches do not require the assumption of a hierarchical bifurcating framework and therefore may be capable of dealing with more complex biological patterns and phenomena [19, 20, 21]. Networks are generally less precise in their ability to reconstruct the divergence points of different groups within a gene family, however, they may be able to capture additional insight into function evolution and divergence using information which might be lost in phylogenetic reconstructions.

In this study we compare the information gained from conventional phylogenetic analysis and a network-based approach using a well characterized subfamily of floral transcription factors, the AGAMOUS floral genes. The AGAMOUS gene subfamily comprises MADS-box transcription factors and is involved in important aspects of flower and fruit development [22]. Among angiosperms (flowering plants), the AGAMOUS subfamily is traditionally divided into the C and D lineages. C lineage genes include the closest relatives of the *Arabidopsis thaliana* AGAMOUS (AG) gene [23, 24] in all angiosperm, as well as close relatives of SHATTERPROOF (SHP) gene, present exclusively in core eudicots.

On the other hand, the D lineage includes angiosperm SEEDSTICK (STK) genes [25, 26]. The C/D split likely occurred after the split between gymnosperms and angiosperms. Thus, gymnosperms usually carry a single gene crop from the AGAMOUS subfamily. While D lineage genes are usually related to ovule development, C lineage genes have been implicated in stamen and carpel development. Particularly in core eudicots, SHP genes have also been shown to be involved in fruit development and ripening [27, 28, 29, 30].

This gene subfamily has been extensively studied and mutant characterization has provided insights into their functional roles in carpel, ovule and fruit development as well as floral meristem termination. The AGAMOUS subfamily has undergone several instances of duplication followed by neo and subfunctionalization throughout its evolutionary history in angiosperms (as reviewed [26, 31]) and understanding the evolutionary history of this group has proven challenging as a result of low support for deep nodes on the tree.

Here we propose using a network approach that analyzes the community structure of the network solely based on sequence identity. Moreover, the network approach used here does not need to calculate gene correlation based on expression data or assume a scale-free topology [32, 16], which makes the approach used more straightforward and robust. Overall, both the phylogeny and network results showed consistent clustering of the gene families. However our results suggest that the network approach was less affected by sequence divergence. We demonstrate that a combination of both methods can may provide additional insight into evolutionary events and functional divergence within gene families.

## 2 Methods

### 2.1 Sequence search and multiple sequence alignment

C and D lineage AGAMOUS nucleotide sequences were retrieved on Phytozome (https://phytozome.jgi.doe.gov/pz/porand NCBI. Species of origin and accession numbers for each sequence included in this analysis are provided in Table 1.

**Table 1:** List of species and sequence identifiers used in this study. Genes retrieved from NCBI (genes with * were retrieved from Phytozome)

| Gymnosperms |  |  |  |
| --- | --- | --- | --- |
| Gene name | Species name | Clade name | ACCESSION or GeneID |
| GinbiMADS5 | *Ginkgo biloba* | Ginkgoaceae | GU563899 |
| PiabDAL2 | *Picea abies* | Pinaceae | X79280.1 |
| PiradAG | *Pinus radiata* | Pinaceae | AF023615 |
| TbaccAG | *Taxus baccata* | Taxaceae | JF519754 |
| CryjaMADS4 | *Cryptomeria japonica* | Taxodiaceae | HM177453 |
| Basal Angiosperms |  |  |  |
| Gene name | Species name | Clade name | ACCESSION or GeneID |
| ShenAG | *Saruma henryi* | Aristolochiaceae | AY464101 |
| ChlspiSTK | *Chloranthus spicatus* | Chloranthaceae | AY464099 |

| PeamAG1 | *Persea americana* | Lauraceae | DQ398021 |
| --- | --- | --- | --- |
| PeamAG2 | *Persea americana* | Lauraceae | DQ398022 |
| MafiAG1 | *Magnolia figo* | Magnoliaceae | JQ326236 |
| MapreSTK | *Magnolia praecossisima* | Magnoliaceae | AB050653 |
| MialAG | *Michelia alba* | Magnoliaceae | JQ326219 |
| **Monocots** |  |  |  |
| Gene name | Species name | Clade name | ACCESSION or GeneID |
| ElguiAG1 | *Elaeis guineensis* | Arecaceae | AY739698 |
| ElguiAG2 | *Elaeis guineensis* | Arecaceae | AY739699 |
| ElguiSTK | *Elaeis guineensis* | Arecaceae | XP_010912706.1 |
| BdiAG1* | *Brachypodium distachyon* | Poaceae | Bradi2g06330.1 |
| BdiAG2* | *Brachypodium distachyon* | Poaceae | Bradi4g40350.1 |
| BdiAG3* | *Brachypodium distachyon* | Poaceae | Bradi2g25090.1 |
| OsMADS3 | *Oryza sativa* | Poaceae | L37528 |
| OsMADS13 | *Oryza sativa* | Poaceae | AF151693 |
| OsMADS21 | *Oryza sativa* | Poaceae | FJ750944 |
| OsMADS58 | *Oryza sativa* | Poaceae | AB232157 |
| SbAG1* | *Sorghum bicolor* | Poaceae | Sb03g002525 |
| SbAG2* | *Sorghum bicolor* | Poaceae | Sb08g006460 or Sobic.008G072900.1 |
| SbAG3* | *Sorghum bicolor* | Poaceae | Sb09g006360 or Sobic.009G075500.3 |
| ZAG2* | *Zea mays* | Poaceae | GRMZM2G160687-T03 |
| ZmSTK2* | *Zea mays* | Poaceae | GRMZM2G471089-T01 |
| ZMM2* | *Zea mays* | Poaceae | GRMZM2G359952-T01 |
| ZAG1* | *Zea mays* | Poaceae | GRMZM2G052890-T01 |
| LaschisAG | *Lacandonia schismatica* | Triuridaceae | GQ214163 |
| LaschisSTK | *Lacandonia schismatica* | Triuridaceae | GQ214164 |
| GongaSTK | *Gongora galeata* | Orchidaceae | AIU94767.1 or KF914206.1 |
| GongaAG | *Gongora galeata* | Orchidaceae | AIU94768.1 |
| HyvilSTK | *Hypoxis villosa* | Hypoxidaceae | AIU94766.1 or KF914205.1 |
| HyvilAG | *Hypoxis villosa* | Hypoxidaceae | AIU94771.1 |
| AspvirSTK | *Asparagus virgatus* | Asparagaceae | AB175825.1 |
| AspvirAG | *Asparagus virgatus* | Asparagaceae | BAD18011.1 |
| **Basal Eudicots** |  |  |  |
| Gene name | Species name | Clade name | ACCESSION or GeneID |
| BgilAG | *Berberis gilgiana* | Berberidaceae | AY464106 |
| EupleAG1 | *Euptelea pleiosperma* | Eupteleaceae | GU357452 |
| EupleAG2 | *Euptelea pleiosperma* | Eupteleaceae | GU357453 |
| AkquiAG | *Akebia quinata* | Lardizabalaceae | AY464107 |
| AktriAG | *Akebia trifoliata* | Lardizabalaceae | AY627635 |
| AktriSTK | *Akebia trifoliata* | Lardizabalaceae | AY627629 |
| HogrAG1 | *Holboellia grandiflora* | Lardizabalaceae | JQ806406 |
| HogrAG2 | *Holboellia grandiflora* | Lardizabalaceae | JQ806407 |
| EscaAG1 | *Eschscholzia californica* | Papaveraceae | DQ088996 |
| EscaAG2 | *Eschscholzia californica* | Papaveraceae | DQ088997 |
| EscaSTK | *Eschscholzia californica* | Papaveraceae | DQ088998 |
| AqAG1* | *Aquilegia coerulea* | Ranunculaceae | Aquca-136-00009.1 |
| AqAG2* | *Aquilegia coerulea* | Ranunculaceae | Aquca-022-00039.1 |
| AqAGL11* | *Aquilegia coerulea* | Ranunculaceae | Aquca-136-00010.1 |
| ThathAG1 | *Thalictrum thalictroides* | Ranunculaceae | JN887118 |
| ThathAG2 | *Thalictrum thalictroides* | Ranunculaceae | AY867879 |
| MedilSTK | *Meliosma dilleniifolia* | Sabiaceae | AY464105 |
| **Core Eudicots** |  |  |  |
| Gene name | Species name | Clade name | ACCESSION or GeneID |
| AlyrAG* | *Arabidopsis lyrata* | Brassicaceae | 946287 |
| AlyrSHP1* | *Arabidopsis lyrata* | Brassicaceae | 486333 |
| AlyrSHP2* | *Arabidopsis lyrata* | Brassicaceae | 321962 |
| AlyrSTK* | *Arabidopsis lyrata* | Brassicaceae | 489841 |
| ATAG | *Arabidopsis thaliana* | Brassicaceae | AT4G18960 |
| ATSHP1 | *Arabidopsis thaliana* | Brassicaceae | AT3G58780 |
| ATSHP2 | *Arabidopsis thaliana* | Brassicaceae | AT2G42830 |
| ATSTK | *Arabidopsis thaliana* | Brassicaceae | AT4G09960 or NM_001203767.1 |
| BraAG* | *Brassica rapa* | Brassicaceae | Brara.K01743.1 |
| BraSHP1* | *Brassica rapa* | Brassicaceae | Brara.G01817.1 |
| BraSHP2* | *Brassica rapa* | Brassicaceae | Brara.E00310.1 |
| BraSTK* | *Brassica rapa* | Brassicaceae | Brara.C02624.1 |
| CaruAG* | *Capsella rubella* | Brassicaceae | Carubv10005558m |
| CaruSHP1* | *Capsella rubella* | Brassicaceae | Carubv10019520m |
| CaruSHP2* | *Capsella rubella* | Brassicaceae | Carubv10025002m |
| CaruSTK* | *Capsella rubella* | Brassicaceae | Carubv10003771 |
| ThhAG1* | *Thellungiella halophila* | Brassicaceae | Thhalv10026567m |
| ThhSHP1* | *Thellungiella halophila* | Brassicaceae | Thhalv10006196 |
| ThhSHP2* | *Thellungiella halophila* | Brassicaceae | Thhalv10017047 |
| ThhSTK* | *Thellungiella halophila* | Brassicaceae | Thhalv10028938 |
| CapaSHP* | *Carica papaya* | Caricaceae | Evm. TU supercontig_50.73 |
| MetrAG* | *Medicago truncatula* | Fabaceae | Medtr8g087860.1 |
| MetrSHP | *Medicago truncatula* | Fabaceae | JX308825 |
| MetrSTK* | *Medicago truncatula* | Fabaceae | Medtr 3g 005530.1 |
| GoraAG1* | *Gossypium raimondii* | Malvaceae | Gorai.N017200.1 |
| GoraAG2* | *Gossypium raimondii* | Malvaceae | Gorai.011G035500.1 |
| GoraSHP* | *Gossypium raimondii* | Malvaceae | Gorai012G042600.1 |
| GoraSTK1* | *Gossypium raimondii* | Malvaceae | Gorai.009G265100.1 |
| GoraSTK2* | *Gossypium raimondii* | Malvaceae | Gorai.009G288000.1 |
| ThecAG* | *Theobroma cacao* | Malvaceae | Thecc1E6029596t1 |
| ThecSHP* | *Theobroma cacao* | Malvaceae | Thecc1EG001841t1 |
| ThecSTK* | *Theobroma cacao* | Malvaceae | Thecc1EG036541t1 |
| MguAG* | *Mimulus guttatus* | Phrymaceae | Mgv1a012605 or Migut.M00986.1 |
| MguSTK* | *Mimulus guttatus* | Phrymaceae | Mgv1a013047m or Migut.C01334.1 |
| PotriAG* | *Populus trichocarpa* | Salicaceae | Potri.011G075800.1 |
| PotriSTK* | *Populus trichocarpa* | Salicaceae | Potri.013G104900.1 |
| PotriSTK2* | *Populus trichocarpa* | Salicaceae | Potri.019G077200.1 |
| TAG | *Solanum lycopersicum* | Solanaceae | L26295.1 |
| TSHP | *Solanum lycopersicum* | Solanaceae | AY098735 |
| TSTK | *Solanum lycopersicum* | Solanaceae | NM_001247265.2 |
| ViviSHP* | *Vitis vinifera* | Vitaceae | GSVIVG01000802001 |
| ViviAG* | *Vitis vinifera* | Vitaceae | GSVIVT01021303001 |

A multiple sequence alignment was performed using the ClustalW [33] alignment tool within Geneious^^®^^ v7.0.4 [34], based on translated nucleotides. Manual curation of the multiple sequence alignment was performed using a codon preserving approach and taking into account domains and motifs previously described in the literature [25]. Unalignable regions were removed prior to further analysis. The final multiple sequence alignment included 549 nucleotides. jModelTest 2.1.1 [35] was used to estimate the best-fit evolutionary model of nucleotide evolution.

### 2.2 Phylogenetic analysis

Maximum likelihood analysis was performed using PhyML 3.0 (http://www.atgc-montpellier.fr/phyml/, [36] [37]) with the TN93 model [38] a gamma distribution parameter of 1.107. Bootstrap support was calculated based on 100 iterations. The most likely tree was computed based on the PhyML estimated parameters: transition/transversion ratio for purines of 2.541, transition/transversion ratio for pyrimidines of 4.342, and nucleotides frequencies of f(A)= 0.33406, f(C)= 0.20359, f(G)= 0.24537, f(T)= 0.21698. Bootstrap support was calculated based on 100 replicates.

### 2.3 Obtaining identity matrix

A pairwise distance matrix, based on a multiple sequence alignment of the 93 sequences was calculated using MEGA7, disregarding nucleotide positions represented by gaps in some species, resulting in a final set of 372 informative positions in the final filtered dataset [39]. The number of base substitutions per site between sequences was calculated using the Maximum Composite Likelihood model [40].

### 2.4 Network analysis

Once the gene identity matrix was generated, a set of 101 networks were created based on the identity threshold between sequence pairs (1 network for each index, 0% through 100%), which is represented by the parameter σ. In each network, each nucleotide sequence is represented by a single node. Two nodes (say *i* and *j*) are considered connected if the identity index is greater than a σ. The networks were represented in the format of an adjacency matrix *M(*σ), where the matrix elements *M*_*ij*_ (pairs of sequences) were either 1, if they were connected, or 0, if they were not connected [41]. Then, neighborhood matrices 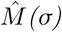 were built for each one of the *M(*σ) [42, 43]. Each element 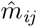 from 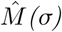 represent the number of steps in the shortest path connecting two nodes *i* and *j*. Whenever two nodes are not connected and belong in different clusters, 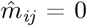. A neighborhood matrix shows the number of edges connecting two nodes in the network. The neighborhood matrices were later used to calculate the network distance δ(σ,σ*+*iJσ) between the pairs of successive networks (in this case Δσ = 1), in order to find the network with the most meaningful biological information, as previously described [41].

GePhi was used to visualize and further interrogate the networks [44]. The modularity calculation from GePhi, based on [45] and resolution from [46] was used to classify individual nodes into communities.

## 3 Results

### 3.1 Phylogenetic Analysis

The Maximum Likelihood phylogeny of AGAMOUS genes presented in Figure 1 is consistent with the topology previously published studies of the AGAMOUS gene family [25, 26, 31]. The most likely tree had a log likelihood score of −20654.546986.

**Figure 1:**
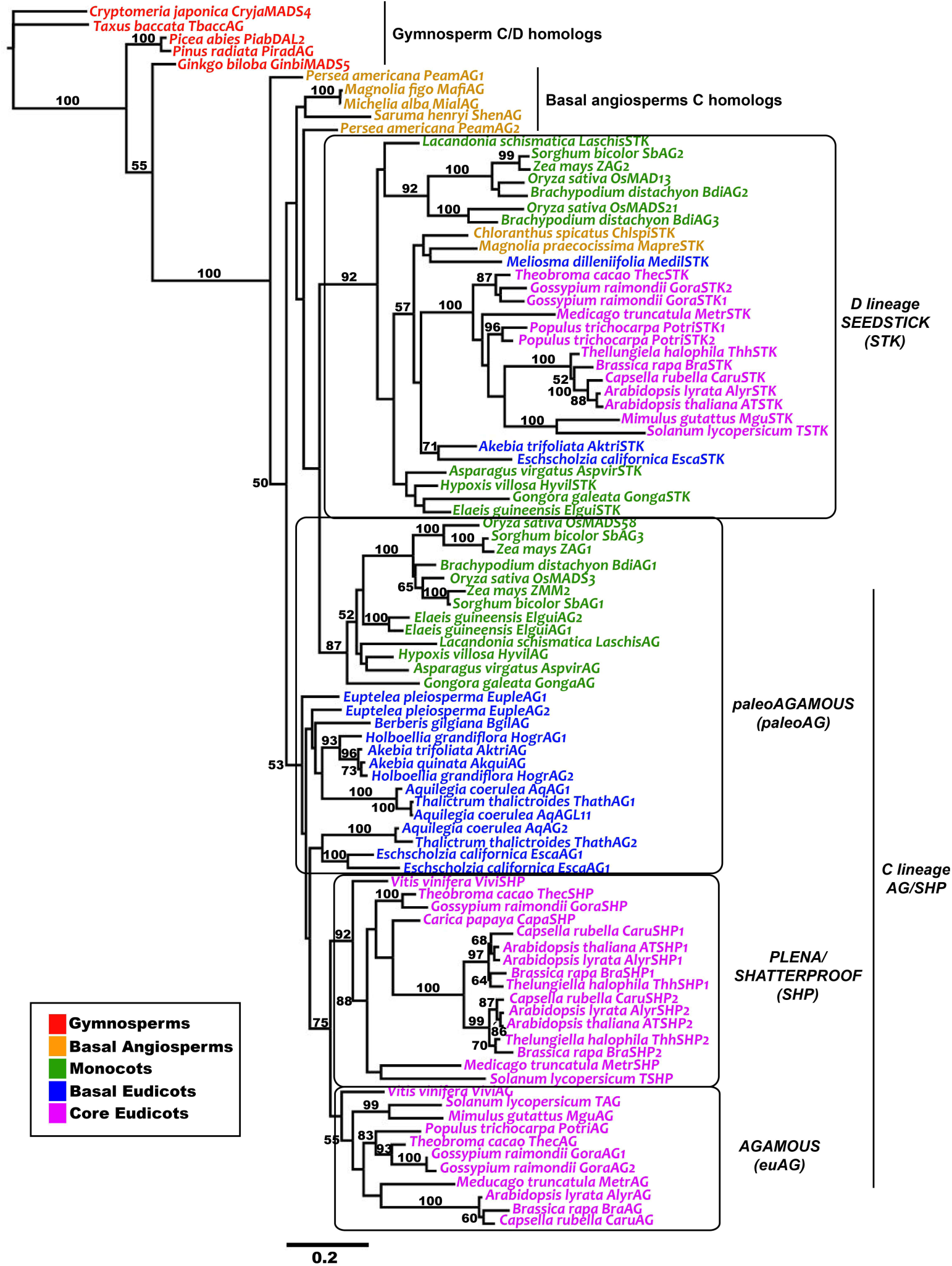
Phylogenetic tree of the AGAMOUS family genes. Main functional groups are highlighted in black boxes along the tree.

Gymnosperm AGAMOUS genes (here termed C/D homologs) form a paraphyly at the base of the unrooted tree. An initial duplication event separates C and D lineage angiosperm genes, and likely occurred in the common ancestor of angiosperms. Basal angiosperm C lineage homologs, although clustering with D lineage genes, exhibit expression patterns, and likely function, similar to that of core eudicot C lineage genes. D lineage genes form a monophyletic clade that includes all other angiosperm species included in this study.

Monocot D lineage genes appear as a paraphyly at the base of the D lineage clade, however the relationships among D lineage genes otherwise are largely consistent with known species relationships. The relationships of C lineage genes is more convoluted. The base of this subtree is a polyphyly including monocot, basal eudicot and core eudicot genes. At the base of the core eudicots, a second duplication event resulted in the split of the AGAMOUS and PLENA/SHATTERPROOF (SHP) lineages. A third duplication, likely at the base of the Brassicales, resulted in two copies of SHP genes in this group (SHP 1 and SHP2) (Figure 1).

Basal angiosperm C lineage genes form a group that diverges after the gymnosperm C/D lineage, but before the angiosperm C/D lineage split. The artificial polyphyletic group of the paleoAGAMOUS include monocot and basal eudicot sequences. While the basal eudicot group with other core eudicot AGAMOUS genes, monocot paleoAGAMOUS genes share a most recent common ancestor with D lineage genes. It is important to notice, however, that the low branch support in many areas of the AGAMOUS gene tree poses challenges to the interpretation of the evolutionary relationship between clades.

### 3.2 Network Analysis

The network distance graph showed its highest peak at 75% identity, which means that the network generated at that peak is the most distant from the others (Figure 2A). Also, it means that the network presents a clear community structure with relevant evolutionary information. Despite the fact that the network with the biggest distance was obtained at 75% identity, the community structure was already too fragmented to answer questions about the evolution of the gene families analyzed in the phylogeny (Figure S1A). A similar situation occurred in [18], and the problem was solved by analyzing other networks in different peaks. Here we attempted to solve this problem by analyzing the network at 51% in order to find the last network where all sequences were connected. However, it was not possible to see a clear community structure in this network due to the high degree of connectivity between nodes (Figure S1B). Finally, in this study we focused mainly on the network obtained at the identity index 67, which meant that two sequences had to have an identity value of 67% or higher to be connected. The choice of the network threshold index was based on the fact that all sequences in this study were connected, with exception of the outgroup sequences.

**Figure 2:**
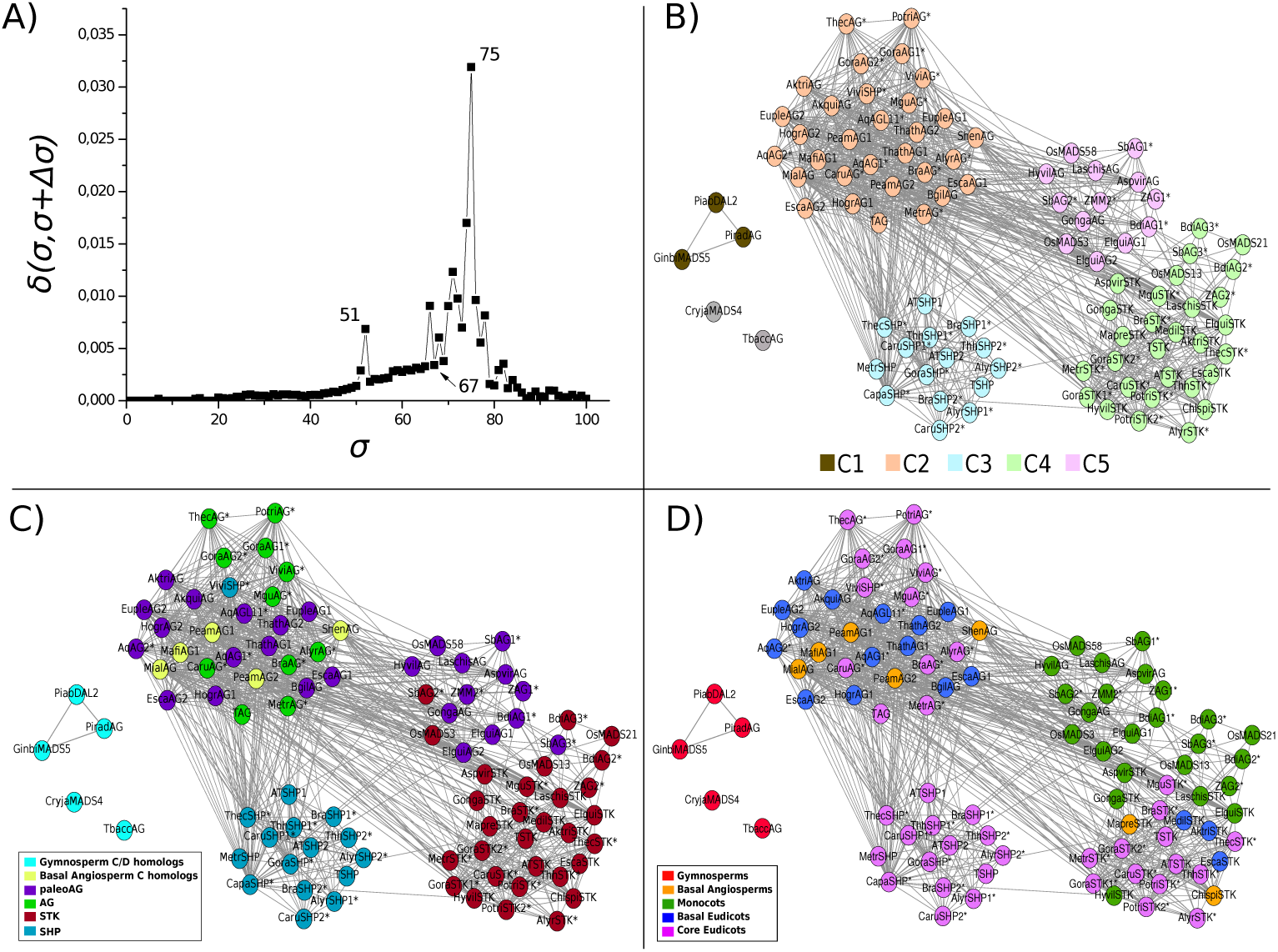
A) Network distance graph based on the *δ(σ,σ+Δσ)* distance. The values for the analyzed networks obtained at 51, 67 and 75% are marked. B) Network obtained at 67% identity. Nodes are colored based on the community they belong to (C1-C5), as result of the modularity algorithm (see methods). The sequences that do not belong to any community are represented as gray nodes. C) Network obtained at 67% identity, colored based on gene function. D) Network obtained at 67% identity colored based on species phylogenetic placement.

After applying the modularity calculation (see methods) in the 67% network, it was possible to see the emergence of the community structure of the network, containing five communities (C1-C5) (Figure 2B). Each one of the communities mainly cluster genes that have similar functions. In C1 3 out of the 5 nodes from Gymnosperm C/D homologs are connected. Even though the 5 nodes are not connected, this result was expected due to the fact that they are part of the most distant outgroup sequences as seen in Figure 1. In C2, on the other hand, the functions of the nodes are related to *AG, paleoAG* and basal angiosperm C homologs. This might suggest that the basal angiosperm C homologs have retained a function very similar to the *AGAMOUS* genes. In C3 the *SHP* genes are clustered together, but in a different community of the *AG* genes, also suggesting functional divergence. The genes clustered in C4 belong to the monocots *paleoAG*. This result might suggest that monocot *paleoAG* genes are evolving under different evolutionary forces than the *paleoAG* and *AG* lineages.

Lastly, C5 comprises the *STK* genes. Even though the communities were mostly composed by genes with similar functions, three genes exhibited unexpected placements. For instance, the *SHP* gene from *Vitis vinifera* (*ViviSHP*) clustered with other AG genes in C2, instead of with other SHP genes in C3. Similarly, *Sorghum bicolor SbAG2*, a *STK* gene, clusted in C5, instead of the expected C4, while *Sorghum bicolor SbAG3*, a *paleoAG* gene, clusted in C4, instead of the expected C5. Finally, we can notice that the grouping obtained by both methods were consistent with one another by comparing figure 2D and figure 1.

Both the phylogenetic and network based analyses returned largely consistent sets of gene clusters. However the grouping of monocots *paleoAG* sequences in a separate cluster (C5) than other C homologs from basal angiosperm, basal eudicot and eudicot sequences (jointly clustered in C2) in the network based analysis suggest two testable hypothesis: (i) monocot sequences are undergoing different and independent evolutionary processes when compared to other non-monocot *AG* homologs, and (ii) non-monocot *AG* sequences are clustered with *euAG* genes due to conservation of function.

## 4 Discussion

The use of phylogenetic methods to study gene family evolution has provided vast increases in the understanding of molecular evolution, and the utility of these methods for reconstructing ancestral relationships remains unparalleled. However, in many cases complex evolutionary processes including neofunctionalization, repeated co-option into new biological roles, as has occurred in independent origins of C4 photosynthesis [47], high birth/death gene families, and reciprocal gene loss following gene or genome duplication, reconstructing phylogenetic relationships may not be the most effective method for identifying genes with equivalent functional roles. Among the contributions of a network approach to gene family studies is the interpretation of the relationships among gene sequences that are not limited to a bifurcating pattern, which is often the case in a phylogenetic framework. A network approach allows for the emergence of patterns that are not seen otherwise. Here we propose the the use of network-based approach which has complementary sets of strengths and weaknesses to conventional phylogenetic methods and tested the contributions of these methods using data from the well characterized AGAMOUS family of floral transcription factors.

In agreement with the literature [48, 26], the network based analysis recovered clusters of *paeloAG* and *AG* genes from basal angiosperms, basal eudicots and core eudicots, potentially indicating conserved functional roles for the genes included in these clusters despite sequence divergence. In contrast, the position of the basal angiosperms C lineage in the phylogenetic tree lead to uncertain interpretations of conserved or divergent function with respect to the D lineage. The network based approach also separated the textitSTK and *paleoAG* genes within the monocot lineage, despite the close phylogenetic relatedness of these two gene clades, consistent with reports of distinct functional roles for these two sets of genes in monocots [49, 50]. For instance, *paleoAG* gene from maize have undergone a duplication event in the common ancestor of maize, wheat and rice [25] which lead to subfunctionalization of these genes, that perform functions still related to, but different from *Arabidopsis AG* [51]. A similar process also occurred in rice [52]. These differences may the the reason the monocot *paleoAG* clustered together in the network, but in a different community than the remaining *AG* gene sequences. Moreover, genetic networks of the inflorescence meristems can vary a lot between grasses and eudicots, since several changes in these regulatory networks are either only present in grasses, or perform a different function in eudicots [53].

However, network-based approaches to studying gene families bring with them their own set of limitations. Some of these are inherent to the particular methodology used here, while others are a result of the relatively immaturity of statistical and software tools for applying these methods to the analysis of gene family evolution. For example, a range of statistical methods are widely available for estimating the level of support for individual branches/clades within a given phylogeny, such as jacknife, bootstrap and posterior probabilities [54, 55]. In contrast, methods for calculation of cluster support in a biological context far less mature, at least for the implementation employed here. The use of sequence identity as a measure of distance, while computationally tractable, also means discarding a great deal of information on the frequency of different types of substitutions at both the nucleotide and amino acid level which can be incorporated into many modern phylogenetic algorithms [56].

Figure 3 summarizes the contributions and relative strengths and weaknesses of phylogenetic and network based approaches to the study of gene family evolution. We propose that the combination of both methods can provide more assessment of both functional and historical relationships between sequences than either approach alone.

**Figure 3:**
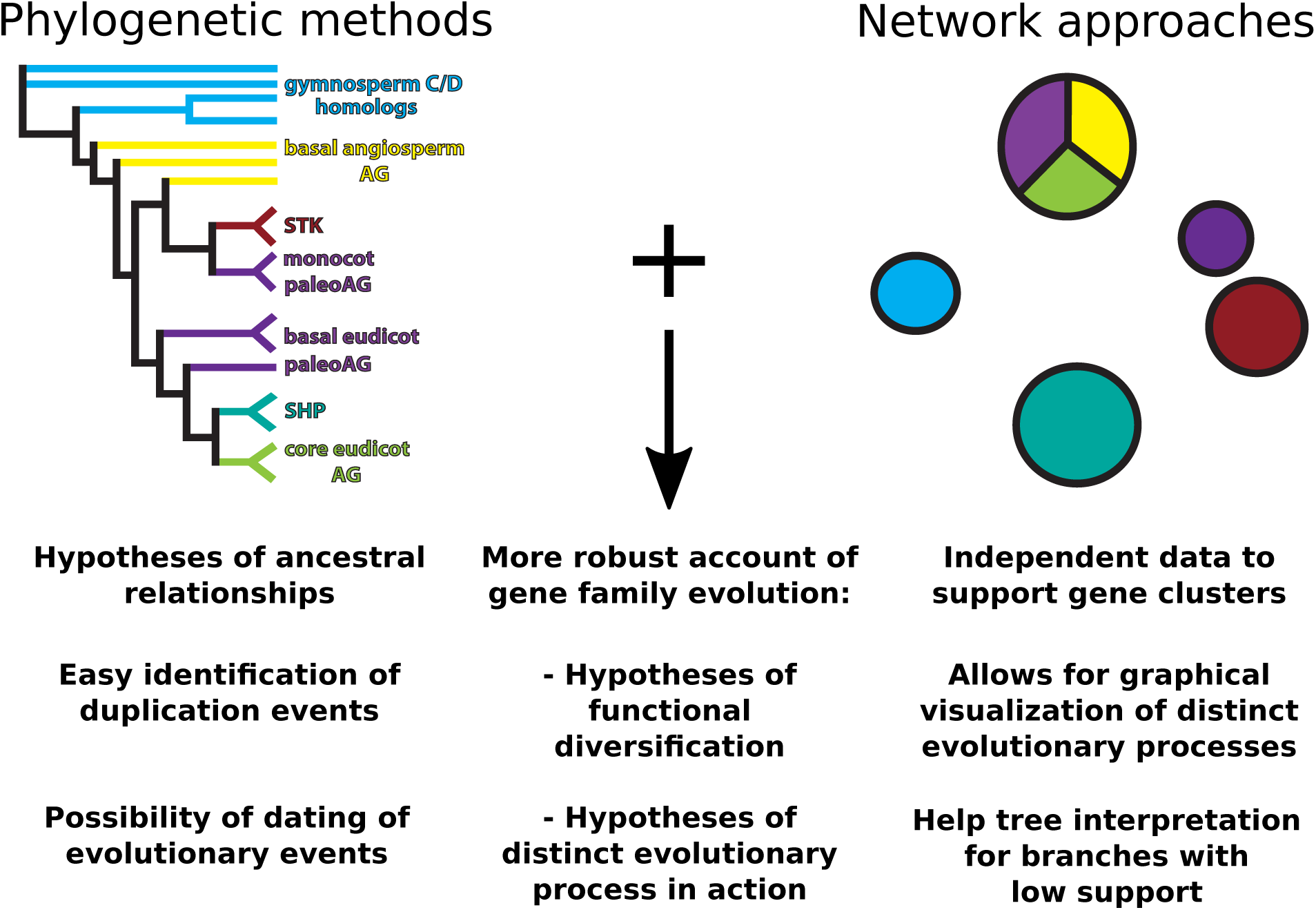
Schematic diagram of results based on phylogenetic (left) and network (right) analyses. Potential contributions of each approach, as well as benefits steaming from the combination of both methods are described below the diagrams.

## 5 Conclusions

Investigating the contributions of a particular network-based approach to the study of the evolution of a well-known family of transcription factor genes involved in floral development supports the idea that network-based approaches, when used in conjunction with phylogenetic methods, can be used to improve our understanding of functional conservation or divergence within gene family evolution. The network-based analysis of gene families used here currently lacks the robust ecosystem of computational tools and statistical approaches developed for phylogenetic analysis, however it can already provide an independent assessment of relationship structures which can aid in the interpretation of phylogenetic data, especially in areas of the tree exhibiting low branch support. In particular, network analysis can be used to generate testable hypotheses regarding the conservation or divergence of gene function in cases of potential subfunctionalization or neofunctionalization. In combination, we believe these methods provide a robust framework that expands the power of gene family evolution studies.

## 6 Acknowledgements

This work was supported by CAPES for fellowship (BJT 069/2013), and Instituto Nacional de Ciência e Tecnologia em Estudos Interdisciplinares e Transdisciplinares em Ecologia e Evolução (INCT-INTREE, 465767/2014-1) to AMRA, a CNPq Science Without Borders scholarship (214038/2014-9) to DSC and a Robert B. Daugherty Water for Food Institute research support award to JCS.

## 7 Supplementary information

**Figure S1:**
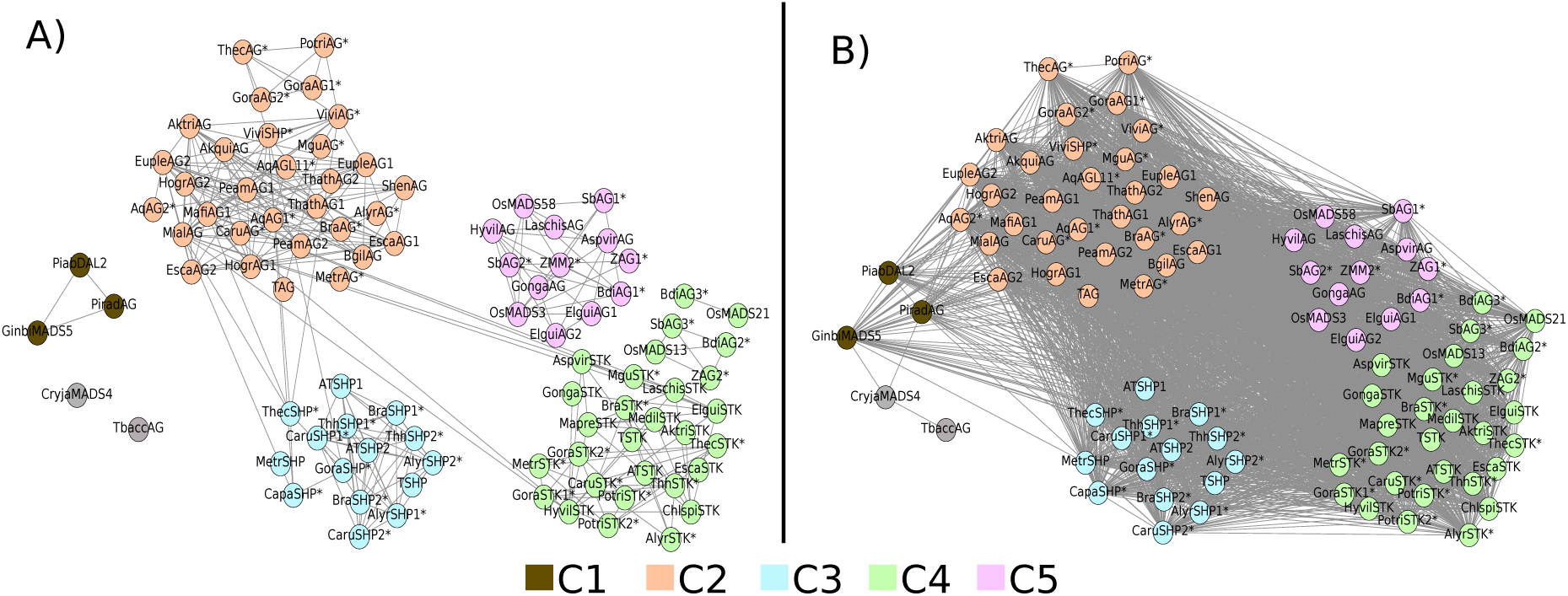
A) Network obtained at 75% identity. Nodes are colored based on the community they belong to in the 67% network, in order to show some community resolution, to highlight the fragmentation of the community structure obtained at 67%. B) Network obtained at 51% identity, also colored based on the community they belong to in the 67% network, despite the fact that the high number of connections did not allow the emergence of community structures. The sequences that do not belong to any community are represented by gray nodes.

